# Enhanced Computational and Experimental Approaches for Comparative Analysis of the Human Mycobiome

**DOI:** 10.1101/2025.02.09.637295

**Authors:** Neelu Begum, Sunjae Lee, Mathieu Almeida, Lars Engstrand, S. Vishal C. Patel, Mathias Uhlen, Dusko Ehrlich, Saeed Shoiae, David L. Moyes

## Abstract

The mycobiome is now recognized as a critical part of the human microbial ecosystem, playing a significant role in both health and disease. Mycobiome research is currently in its infancy and relies on fungal-specific primers with challenges in bioinformatics to accurately determine the structure of the mycobiota community. In addition, the majority of computational and experimental methods currently in use have been optimized for bacteria rather than for fungi. Here, we provide a comparative analysis of extraction methodologies for metagenome sequencing and subsequent bioinformatic analysis that will enhance fungal species identification. We utilized cultured mock fungal communities, including both yeast and mold species, to evaluate the efficiency of extraction protocols. We further enhanced computational analysis of the mycobiome, using mock and human metagenome data together with our curated catalogue consisting of 984 fungal genomes specific to the human mycobiome. Application of this optimised workflow for oral and gut samples of healthy individuals detected of novel species from *Puccinia* and *Lentinus* genera among the established presence of *Malassezia*, *Rhizophagus*, *Candida* and *Saccharomyces* genera. In addition, *Enterocytozoon,* was identified specifically in the gut mycobiome. Our pipeline enabled the comprehensive sampling of all fungal genes in a microbial community within a human sample minimising bias, reducing errors and artefacts from amplification, and providing accurate diversity and abundance data for the human mycobiome.

## Introduction

Fungal diseases affect over a billion people every year, with mortality and co-morbidity estimated at around 1.5 million per year^1^. This is being exacerbated by global warming driving environmental adaptation of invasive fungi which are becoming a clinical concern^2^. Fungi have been associated with various human diseases, with shifts in mycobiome composition found in everything from invasive fungal infection to immunomodulation, and more recently driving cancer progression^3–8^. In addition, antifungal resistance is a rising problem with limited anti-fungal drug options in clinical settings and a high degree of similarity of these drugs with agrichemicals used in farming^9^. Investigation of both isolated fungal species and fungal communities (mycobiome) and their interactions with bacteria and host is critical to developing future therapeutic interventions in these infections^10–12^. Developing new approaches to investigating the mycobiome is therefore fundamental to our understanding of how these microbes contribute to human homeostasis and pathology.

To fully explore the mycobiome, it is essential to begin analyzing these communities using shotgun metagenomics approaches. However, there is currently a lack of gold standards for sample preparation and extraction of mycobiome DNA and subsequent analysis to facilitate increasing mapping rates to known genomes and *de novo* assembly of new fungal genomes. Fungal DNA extraction has long been linked to individual species-specific protocols due to phenotypic differences such as melanin in the cell wall and filamentation^13–18^. Thus, the robustness of complete mycobiome analyses is more difficult. Extraction protocols are often labour intensive with heavy chemical use, whilst commercial kits show variation in their ability to extract DNA from different fungal species^19–21^. For example, *Malassezia spp.* have a thicker/tougher cell wall in comparison to other fungal species resulting in problems extracting their DNA efficiently using standard protocols^22^. With increasing recognition of the importance of the mycobiome, there is a need to explore more accurate, large-scale, and efficient methods of sample processing^23^. In addition to the DNA extraction issues, identification of species within the mycobiome is currently predominantly at the genus level, using targeted amplification of the Internal Transcribed Spacer (ITS) region and subsequent sequencing^24^. However, use of polymerase chain reaction (PCR) results in sequence bias of known species, increases the incidence of duplication and has recently been identified to be inaccurate for relative abundance calculation due to duplication of ITS regions in different fungal species^25,26^. Baseline fungal genomics still relies on amplicon-based methods to establish fungal diversity and abundance. Recent advances in high-throughput technologies pioneered by the Human Microbiome Project (HMP)^27^ and METAgenomics of the Human Intestinal Tract (MetaHit)^28^ have allowed for the high-resolution characterisation of the bacterial/archaeal component of the human microbiome. However, the assumption that fungal genes typically make up less than 0.1% of the human microbiome, coupled with the limited number of annotated fungal genomes, has resulted in the underdevelopment of fungal exploration^29^. There are a few exisiting fungal pipelines for metagenomics analysis such as FindFungi^30^ and Human Mycobiome Scan (HMS)^31^. However, these resources are limited for fungal genomes, coverage of the mycobiome and generation of gene counts for further in-depth functional annotations.

Here, two pipelines for mycobiome DNA extraction were benchmarked. We tested two extraction protocols commonly used in microbiome research and developed an improved extraction protocol for mycobiome sampling that has been adapted for improved lysis of human associated fungal cells. To standardise these procedures, we used a mock community comprised of 12 distinct fungal species to benchmark extraction protocols and compared a novel computational pipeline to analyse shotgun metagenomic mycobiome data with existing tools^31–33^. Finally, we demonstrate the strength of the proposed extraction and computational workflow by applying it to saliva and stool samples from healthy individuals. In doing so, we provide a new baseline standard for shotgun metagenomics sequencing of the human mycobiome alongside a tool for the discovery of unculturable fungal species thereby providing a platform for expanding human mycobiome research.

## Methods

### Establishment of fungal mock communities

12 species were included in the mock communities used in this study: 7 *Candida* species *- C. albicans*, *C. dubliniensis*, *C. tropicalis*, *C. glabrata*, *C. auris*, *C. parapsilosis*, and *C. krusei* (also known as *P. kudriavzevii*); other common yeast human fungal pathogens - *Cryptococcus neoformans* and *Cryptococcus gatti,* two mold species *Aspergillus fumigatus,* and *A. brasiliensis,* and the common commensal yeast species *Saccharomyces cerevisiae* (**Supplementary table 1**). For optimal growth of all 12 fungal species, cultures were grown at 25°C for 3 days in Sabouraud-dextrose broth (SAB) using an orbital shaking incubator at 95rpm (Sabouraud-dextrose broth, Oxoid UK; SI500 benchtop, Cole-Parmer).

### Healthy Sample collection

Three healthy individual saliva and faecal samples were collected in a previous study^34^.

### DNA extraction and quantification

DNA from the mock community samples, healthy saliva and faecal samples was extracted using two protocols; DNeasy PowerSoil Pro Kit (Qiagen) and chemical extraction procedure developed at MetaGenoPoliS (MGPS)^35^. All extractions were performed in triplicate according to previous protocols with adaptations. Throughout this paper we refer to the adapted PowerSoil Pro Kit as PKS and chemical extraction protocol as MGPS. An initial lysis adaption was added to bot h protocols to optimise for extraction from fungal species. This involved addition of three protocols: (i) using 0.5 mm diameter yttria-stabilized zirconium oxide beads (Lysing Matrix Y, MP Biomedicals) (ii) heating the sample for 10 min at 70 °C and (iii) performing a bead-beating at 6.5m/s every 30secs with 3 min intervals ice repeated 6 times leading to a total of 3 min beat beating (FastPrep-24, MPBio).

DNA quantification was carried out using the dsDNA HS Assay (Invitrogen Qubit®, ThermoFisher) with Qubit® standards. DNA quality was assessed using gel electrophoresis. Primer sequences used for quality control can be found in Supplementary table 2. Quantitative PCR was performed on a Rotor-Gene 6000 (Qiagen, UK) with cycle information found in Supplementary table 3.

### Mock community evaluation

The mock community was assessed using three protocols: (i) detection of individual species using qPCR in mock community sample, (ii) growth rate measured of individual and mock and (iii) determine dry cell weigh (DCW). The species in the mock communities were detected using species-specific primers in a qPCR assay (**Supplementary table 2; Methods**). The cycle threshold (Ct) values of individual strain and mock community DNA reactions were used to calculate relative abundances. Inverse proportion of Ct values on the graph was calculated by dividing per species against total mock community Ct value. The growth rate was measured, by determining the absorbance at 600 nm and DCW for each species and the mock communities to determine microbial load and confirm growth of individual strains within the mock communities.

### DNA Sequencing

The initial quality control assessment and subsequent sequencing was performed by National Genomics Infrastructure (NGI) in Sweden. The Illumina TruSeq PCR-free library preparation kit (Illumina, USA) was used to generate sequence libraries for DNA with 350 bp average fragment size. DNA sequencing was subsequently performed on a NovaSeq 6000 (Illumina, USA) generating a minimum sequencing depth of 20 million reads per sample after filtering reads for paired end reads for mock and human samples^34^.

### Mapping to the fungal reference catalogue

The fungal catalogue used in this study was previously generated ^36^ and is available at https://entrepot.recherche.data.gouv.fr/dataset.xhtml?persistentId=doi:10.15454/WQ4UTV. The catalogue consists of 2,440,644 fungal genes derived from 984 fungal genomes. The reference genes were assembled into the fungal catalogue and can be utilised via the Meteor software suit^37^ with the parameters of *-p* (cpu count) *10*, -*trim3* option, read_length >35 and *-k* (aligns per read) based on match count for Bowtie2. Non-fungal reads were removed, including human reads (Homo_sapiens_GRCh38; https://www.ncbi.nlm.nih.gov/assembly/GCF_000001405.26/), bacterial reads (hs_10_4_igc2)^37^, animal reads (Bos_taurus_UMDS; https://www.ncbi.nlm.nih.gov/assembly/GCF_000003055.6/) and plant reads (A_thaliana_TAIR10; https://www.ncbi.nlm.nih.gov/assembly/GCF_000001735.3/). The gene count was produced from the mapping scores of samples against the fungal catalogue using Bowtie2 built within Meteor^37^ framework. Gene count was normalised based on reads per kilobase of exon per million mapped (RPKM) based on the *momr* package (version 1.1) in R. Downstream analysis cut-off of 62% was established and relative abundance was achieved using an R script. The pipeline has been automated into a Nextflow pipeline that can be found at https://github.com/sysbiomelab.

### Evaluation of fungal catalogue outcomes with other pipelines

Human Mycobiome Scan (HMS) version 1 is publicly available and was downloaded from https://sourceforge.net/projects/hmscan. Sequences were submitted for profiling using fungi_LITE that included 66 full genomes. Metaphlan3.0 is another bioinformatic pipeline used to speciate our sequences using ‘mpa_v30_CHOCOPhlAn_201901’ database, this included 500 eukaryotes genomes. Installation and download of Metaphlan3.0 was available at https://huttenhower.sph.harvard.edu/metaphlan/ and https://github.com/biobakery/biobakery/wiki/metaphlan3 [accessed: September 2021].

### Evaluation of fungal catalogue mappings against 18S markers

Unmapped reads and unique reads were assessed using SAMtools and BAMtools from Metagenomics tool (https://www.metagenomics.wiki/tools/short-read/remove-host-sequences). ITS/18S reference dataset markers were blasted against mock community and compared to fungal profiling with our optimised pipeline. Blastn was used for single-best hit against the UNITE database ^38^ to retrieve ITS and 18S genus taxonomic identification against mock community reads. Single-best hit means to receive one hit per gene based on highest percentage identity with e-value 0 to reduce noise. The Pyani package^39^ was used to assess genomic similarity between fungal species.

### Statistics

All statistical analyses were performed using R software v3.6.3 to evaluate means and standard errors for boxplots.

## Results

### Evaluation of mycobiome DNA extraction using fungal mock communities

We began our study by cultivating multiple fungal species in a range of different communities to test and evaluate different DNA extraction workflows. This approach allows for a more precise representation of the mycobiome community within a controlled laboratory setting. Our mock community comprise of 12 different fungal species across 4 genera. To validate that the fungal species were able to co-exist in a community, we determined the growth rate and species composition of the communities (**Figure 1a Methods**). We then used these mock communities to assess the robustness of different extraction protocols. Later, we used the same protocol on human saliva and faecal samples, together with different metagenome mapping pipelines for detection of the fungal species (**Figure 1b; Methods**). We initially tested two different commonly used microbial DNA extraction protocols on a fungal mock community consisting of 12 species (*Candida, Aspergillus, Cryptococcus* and *Saccharomyces* genera (**Figure 1b; Methods; Supplementary table 1**) representing a mycobiome community. We compared an adapted commercial DNeasy PowerSoil Pro Kit (PKS) and the Metagenopolis chemical extraction protocol (MGPS) ^35^ extraction protocols that we optimised for concurrent fungal and bacterial cell lysis. To improve lysis of fungal cells, protocol was adapted with an initial heat treatment at 70°C, lysing matrix Y (MP Bio) and specific mechanical homogeniser protocol (Methods). MGPS extraction yielded ∼330 ng/µl of DNA, whereas the PKS pipeline yielded ∼20 ng/µl (**Figure 1c**). Using DNA fragmentation as representative of DNA quality for shotgun sequencing, the PKS protocol gave an average fragment size of ∼6,289 bp whilst MGPS gave ∼869 bp (**Figure 1d**). Both protocol average fragment sizes were larger than the requirement for NGS sequencing. However, due to the similarity of fungal species genome assembly, large fragment size will provide more accuracy, especially for long read sequencing technologies^40^. Both extraction workflows were validated using qPCR with specific individual primers for the detection of known species in the mock community, showing successful isolation of all fungal species with both extraction processes (**Supplementary Figure 1a**). However, using ITS/ITS2 universal primers, we found the PKS protocol gave lower average cycle threshold (ct) values, compared to the MGPS protocol indicating detection of higher amounts of fungal DNA in the starting reaction (**Supplementary Figure 1b**).

**Figure 1.**
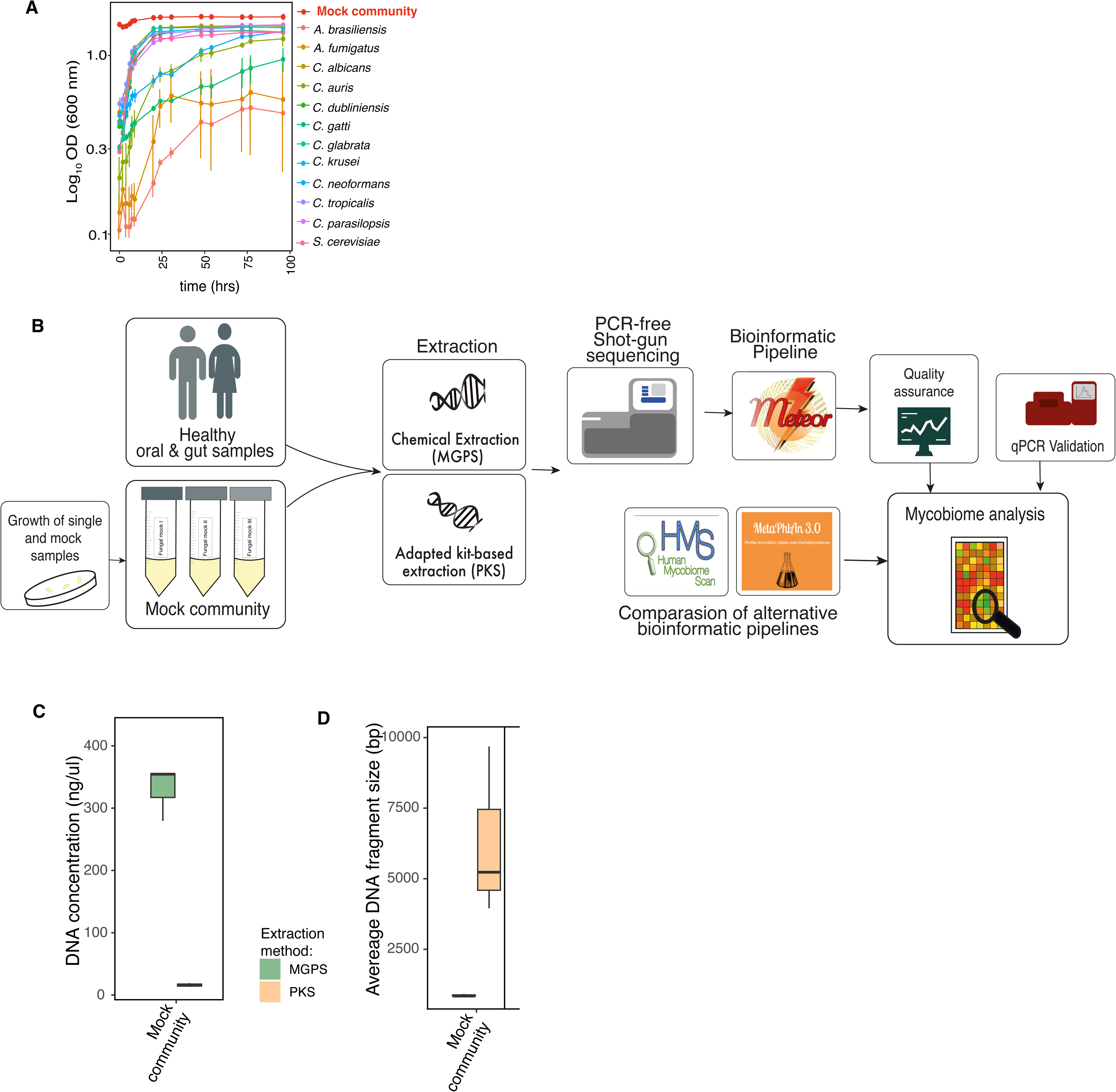
**A)** Individual growth of 12 fungal species was compared to mock community (inclusive of all 12 fungal species) measured over 100hours at absorbance of 600nm (see Methods). Mean OD was plotted with biological triplicate repeat ± SD ymin/ymax. **B)** The workflow of comparative experimental, data generation and analysis to obtain optimal procedure for mycobiome analysis, and results from the in-vitro experiments. **A)** A detailed schematic of workflow performed to establish an optimised DNA extraction and sequencing for the human mycobiome. DNA of mock community and healthy human samples (oral and gut) was extracted using a chemical extraction kit (MGPS) and adapted protocol of Qiagen Powersoil Microbiome Kit (PKS). PCR-free shot-gun metagenome was performed and bioinformatic pipeline was applied to retrieve relative abundance. This was compared with existing bioinformtic tools (HumanMycobiomeScan and MetaPhalAn 3.0). Additional laboratory validation was performed using qPCR. This allowed us to compare most effective method in DNA extraction of human mycobiome profiling. **C)** DNA concentration of mock community retrieved from chemical (MGPS, green) and kit-based (PKS, orange) extraction. Biological triplicates mean ± SD. **D)** Average DNA fragmentation size of mock community measured for MGPS and PKS extraction protocols. Biological repeats mean ± SD.

### Mycobiome analysis using fungal catalogue

Next, to improve the identification and accurate quantitation of fungal species in the mycobiome, extracted DNA from the mock community was sequenced to a depth of minimum 20 million reads. These reads were directly mapped against our fungal non-redundant gene catalogue^36^. After the gene counts were produced for each sample, normalisation was performed to retrieve relative species abundance (**Methods**). In the top 50 abundant fungal species recovered from alignment, there was an accurate recall of 11 of the 12 fungal species present in the mock community (**Figure 2a**). Interestingly, the additional species identified (above the second blue line) were closely related species within the same genus to those in the mock community as these species can be mis-identified. The additional species sequences were compared to the species from the mock community using average nucleotide identity, indicating a high degree of sequence similarity in these closely related species (**Supplementary Figure 2a**).

**Figure 2.**
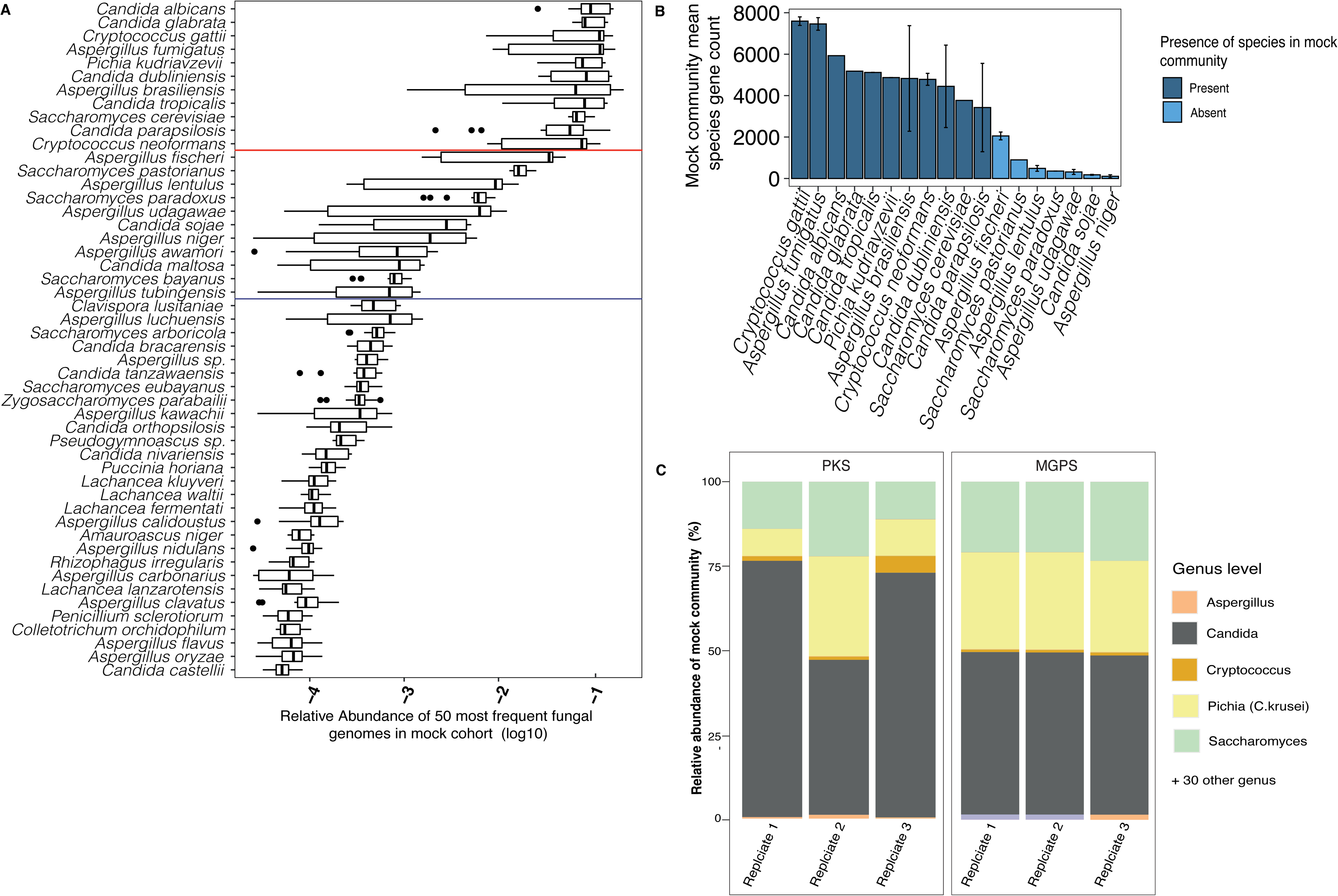
Assessment of extraction and fungal catalogue using mock community **A)** Top 50 relative abundant species identified from mapping to fungal catalogue. Red line indicates top 11 species, and second blue line indicates top 20 species. **B)** Mean gene count of species recalled from mock community from in-house catalogue mapping. Dark blue indicating species present in the mock community and light blue indicates fungal species absent in mock community, indicating inaccuracies. Error bars indicating mean ± SD. **C)** Comparison of relative abundance of mock community genus with MGPS and PKS. Each replicate represented to demonstrate sample variation. To note, *Pichia kudriavzevii* was relabelled as *Candida krusei* as error in identification of isolate and deposit of sequences in NCBI have not been corrected^57^.

The individual replicate DNA samples taken from the mock community with each extraction technique highlighted accurate recall of species. Notably, however, the MGPS extraction pipeline resulted in identification of an additional 30 fungal species that were not present in the culture, although this did not cause a distortion of the relative abundance of mock community (**Figure 2b**). In contrast, the PKS protocol recalled fewer closely related fungal species compared to MGPS, although PKS had more variation in the relative abundance of *Candida* species across the three biological replicates. Assessing the mean raw gene count of the mock community across the replicates of PKS-extracted DNA, we can identify a threshold for accurate cut-off between present species and closely related species (**Figure 2c**). Examining the breadth of coverage of the mock community, we established the sequencing depth of 20 million reads is sufficient to detect fungi on a species level from mycobiome sampling using the revised protocol used in this work(**Supplementary Figure 2b**). The overall alignment of sequences from mock community samples was higher for DNA extracted using the PKS extraction protocol as compared to those extracted with the MGPS protocol with >50% alignment (**Supplementary Figure 2c**). In contrast, sequences from the MGPS pipeline were mapped to more closely related species than the PKS pipeline. This could be due to DNA shearing creating smaller fragment size in MGPS extraction leading to mapping to more closely related species from sequence similarity thus creating increased false positives.

### Comparison of mycobiome mapping using other resources

To evaluate the results from our mycobiome catalogue pipeline analysis relative to other existing pipelines, we compared these results to those generated using other available resources. To do this, the same sequence reads from mock community DNA samples extracted with both pipelines were compared across the different fungal metagenomic pipelines. There is 56% alignment of reads to the fungal catalogue with PKS and with MGPS extracted DNA having an average of 60% of unmapped reads (Supplementary Figure 3a). In addition to our fungal catalogue pipeline, we were able to map reads to the species level using both the HMS and MetaPhlAn 3.0 pipelines (**Figure 3a**). The fungal catalogue pipeline identified 11 out of the 12 species present in the community. In contrast, both the HMS and MetPhlAn 3.0 pipelines identified 6 of the 12 species. Notably, *Candida auris* was not identified by any of the pipelines. For the fungal catalogue pipeline, this was due to non-inclusion of *C. auris* genomes in the catalogue version used here^36^.

**Figure 3.**
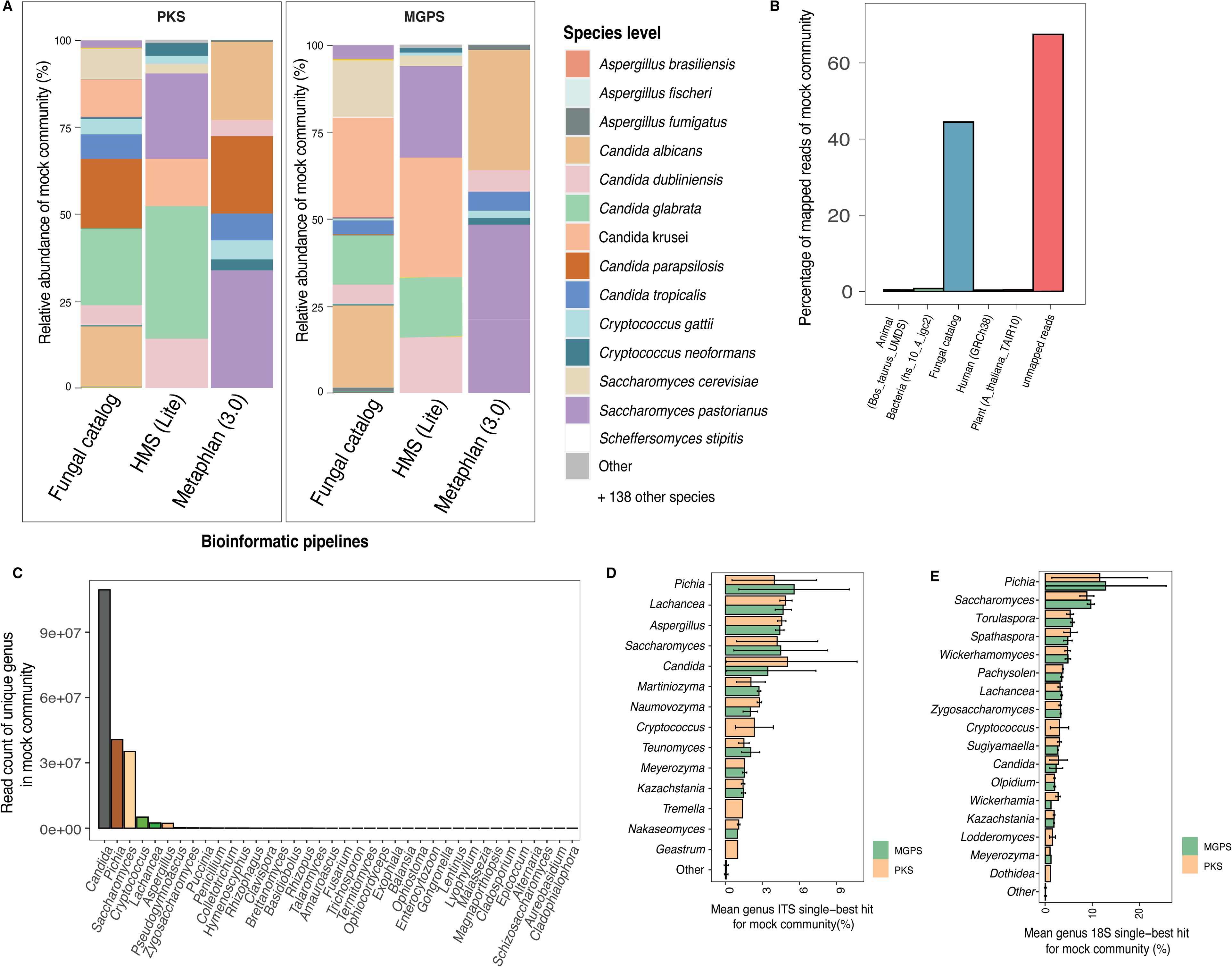
Comparative analysis of bioinformatic pipelines for precision and recall of fungal species. **A)** Comparison of relative abundance of mock community for species level annotation applied across in-house catalogue, Mycobiome Scan (HMS) and MetaPhlAn (version 3.0). Biological triplicates. **B)** Percentage of mapped reads across different reference genomes and catalogues.The fungal catalogue assigned to most reads. There is no sample cross-contamination with animal, bacteria, human or plant. The unmapped reads appear to make up the remainder of the missing reads. **C)** Unique read count of mock community was mapped to determine accuracy of ungal catalogue mapping. The sample presence of *Candida, Saccharomyces, Cryptococcus, Aspergillus* was confirmed. The detection of *Pichia* and *Lachancea* read counts may be attributed to sequence similarity. **D)** Mock community sequence was mapped to ITS library, for quality detection using single best hit to reveal genus occurrence. The bars are represented in colour green for MGPS and orange for PKS. Error bar represented by mean ± SD of biological triplicates. **E)** Single best hit percentage of mock community were mapped to 18S library for detection of genus. Genus non-existing in mock community was mapped at a higher percentage than *Candida, Cryptococcus* and A*spergillus* was not detected. The bars are represented in colour green for MGPS and orange for PKS. Error bar represented by mean ± SD of biological triplicates.

We next performed further evaluation to understand the distribution of mapping of the mock community reads. Mapping against bacteria (hs_10_4_igc2), animal (Bos_taurus_UMDS), human (Homo_sapiens_GRCh38) and plant (A_thaliana_TAIR10) reference genome catalogues showed no hits, indicating that the reads were either fungal species or unassigned, demonstrating efficient processing in the bioinformatic pipeline using the fungal catalogue (**Figure 3b; Methods**). The unique reads from the mock community sequences were captured to determine the accuracy of genera mapping. *Candida*, *Saccharomyces, Pichia and Cryptococcus* accounted for majority of genera detected (**Figure 3c**). As mentioned earlier, the *Pichia* genus has now been reassigned to the *Candida* genus due to sequence similarity ^19^. Other examples of taxonomy rearrangement due to sequence similarity of closely related species include the *Lachancea* genus that was separated from *Saccharomyces* as a new genus in 2003^21^. This highlights the need for both sequencing and reconsolidation of fungal taxa using NGS^41^. To determine if any of the unassigned reads belong to *Protista* genera, the unmapped reads were mapped against NCBI (**Supplementary Figure 3b**). We found the unassigned reads mapped to other strains of *Candida, Pichia, Cryptococcus* and *Saccharomyces* that weren’t added to the fungal catalogue. This highlights the need to regularly update the fungal catalogue with known species and *de novo* assemblies in the mycobiome. Additionally, the confirmation of missing *C. auris* species in the fungal catalogue and other strains of *C. albicans* not included reduced the overall alignment of mapped reads (**Supplementary Figure 3b**). This indicates that similarity of annotations across the species may highlight genera inaccuracies, meaning that more strains of fungal species need to be added to the fungal catalogue for robustness or else genera sequence similarity may not be sufficient for fungal species differentiation in the future. Genus classification using ITS and 18S using single-best hit was performed on mock community to reveal the relative abundances of genera (**Figure 3d-e**). ITS performed much better than 18S with single-best hit, identifying *Pichia, Aspergillus, Saccharomyces, Candida, Cryptococcus* but mycobiome profiling was masked by non-related species abundancies. In contrast, 18S analysis failed to identify all genera within the mock community (**Figure 3e**).

### Metagenomic analysis of human mycobiome samples

To demonstrate the applicability and reproducibility of the workflow in complex environmental/human microbial communities, the PKS DNA extraction and fungal catalogue workflow was applied to paired saliva and faecal samples from 3 healthy participants. The overall DNA extraction for both oral and gut was high with yields of ∼70-100 ng/µl (**Figure 4a; Supplementary Figure 4a-b**). The DNA was sequenced to a minimum depth of 20 million reads per sample. Mapping the samples indicated we were able to obtain good coverage of fungal annotation of the mapped reads (**Supplementary Figure 4c**). The alignment of the fungal species reads to the catalogue was around 1% (Figure 4b) which could be due to the low abundance of fungal genes reported in the human microbiota^29,42^. For clarification of the effectiveness of shotgun metagenomics, reads were aligned to other non-fungal genomes, revealing >75% mapping to the human genome and ∼14% mapping to our oral bacterial reference genome catalogue for DNA from saliva (Figure 4d). The stool samples revealed >75% annotation to a bacterial catalogue^37^, which is consistent for known constructs of the human microbiota and the low abundance of fungi in these communities^29^.

**Figure 4.**
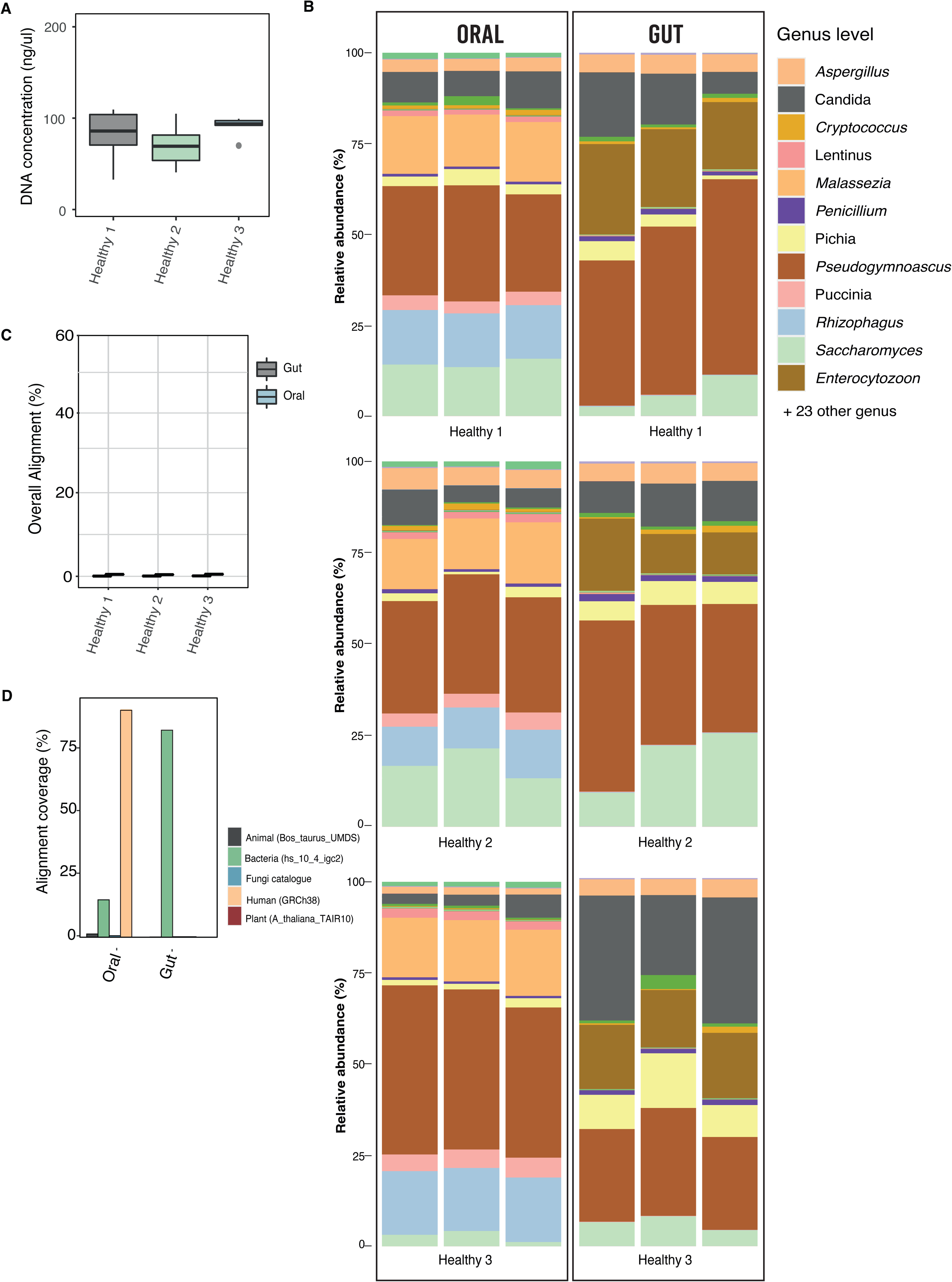
Clinical application of optimised mycobiome pipeline on oral and gut samples. **A)** DNA quantitation of both healthy oral and gut using PKS extraction. Error bar represented by mean ± SD of biological triplicates. **B)** Total alignment of fungal catalogue in healthy oral and gut participates, indicating the lesser presence of mycobiome in human. Oral and gut sample triplicates has been shown for each healthy participant represent differentiation. Error bar represented by mean ± SD. **C)** Genus level relative abundance comparison of healthy oral and gut mycobiome profile. Individual triplicate shown to demonstrate consistency in genus recall. **D)** Read distribution of oral and gut healthy participant samples to animal, human, plant, bacterial and fungal genomes, and catalogue. The oral samples sequences were attributed to majority of human and bacterial catalogue. The gut samples were mapped to bacterial catalogue.

Using the fungal catalogue, the taxonomic profile was characterised showing the presence of the previously identified genera of *Malassezia, Rhizophagus, Candida* and *Saccharomyce*s in gut mycobiome (**Figure 4c**) ^43–47^. In addition, we observed the presence of novel genera including *Puccinia* and *Lentinu*s in the oral mycobiome. Notably, the *Lentinus* genus are club fungi including edible fungi such as shiitake mushrooms, thus its presence in these samples may be due to dietary contamination^48^. Similarly, given their common role as plant pathogens consumption of wheat/whole grains/seeds could be responsible for the presence of the *Puccinia* genus^49^. Interestingly, although previous studies have highlighted a dominance of *Saccharomyces* and *Candida* spp in the mycobiome^50–53^, these genera are not as dominant as previously determined with this pipeline.

## Discussion

The current standard use of ITS amplicon sequencing for all mycobiome analysis has recently been called into question as it introduces errors in the relative abundances of fungal species due to hybridisation from point mutation, introgression from recombination and gene duplication in pseudogenes^54^. In addition, this pipeline suffers from being a purely phylogenetic analysis, providing none of the functional analyses possible with metagenomic analyses that are becoming ever more revealing of host-microbiome interaction mechanisms. Comparative analysis of our samples against ITS/18S database confirms recent reports of multiple copies of ITS region causing skewed assessment of relative abundances^25^. The comparison of the UNITE database and our workflow indicates that our pipeline incorporates the most efficient method and tools for fungal species annotation and abundance calculation, whilst also avoiding the use of OTUs and amplicons.

In this study, we introduce an optimised workflow for joint fungal/bacterial DNA extraction from different samples for subsequent metagenomic analysis. In comparing our pipeline to ITS and 18S databases we highlighted mis-annotations, leading to further doubts regarding the accuracy of ITS sequencing for analysing the mycobiome. In order to mitigate the issues of assessing fungal abundance, we developed a revised DNA extraction protocol with reduced operator variation by using kit based method, and a subsequent bioinformatic analysis pipeline to allow the human mycobiome to be investigated comprehensively. This workflow was designed to meet clinical and research needs for mycobiome compositional analysis but will also allow the functional analysis inherent in shotgun metagenomics to be carried out on fungal communities. Several kit based extraction protocols have been tested in other studies for microbiome DNA analysis^24^ still using amplicon methods, but with the PKS protocol providing the better quality and quantity of DNA required for shotgun metagenome sequencing. However, we must acknowledge these extractions have all been optimised for bacterial microbial DNA. We report PKS is sufficient for shotgun metagenomic profiling of the mycobiome, but the DNA quantity was lower (approximately ∼20ng/µl) than the traditional chemical extraction protocol. On the other hand, PKS provided larger average fragment size, meaning that the PKS extraction gave better DNA starting material sequence alignment, as well as providing longer average fragments for future use in 3^rd^ generation sequencing technologies such as PacBio and Oxford Nanopore. Furthermore, the PKS protocol is significantly quicker meaning shorter handling times, reduced operator variation and easier large scale mycobiome sampling for use in clinical settings. In contrast, the MGPS protocol takes longer (2 days) with a need to make a range of reagents that is time consuming, open to contamination and produces smaller fragment size. However, MGPS provides higher DNA quantity that may prove useful for single species application or deeper sequencing. Here we add a series of further additions to the protocol - the use of larger mechanical beads, a heating step and incubation on ice during homogenisation. These steps facilitate breaking open the tough fungal cell wall whilst also preventing degradation of fungal DNA. PKS has been certified for use in microbial clinical metagenomics^55^ and its use can now be extended to include mycobiome analysis. This PKS extraction protocol can be employed for sample extraction on a large-scale by non-specialist users, allowing for decreased inter-operator variation in DNA yield and quality without the need for an experienced individual and reducing downstream variation both within a study and in comparison to different studies.

The fungal catalogue developed and utilised for the first time here^36^ underwent extensive curation using literature searches to identify fungal species that have been previously seen in humans. This pipeline provided more specific and accurate recall of fungal species as compared to other commonly used pipelines and with specificity to human-related fungal species. It is important to note, however, that accurate curation of this fungal catalogue^35,56^ will involve regular updates and expansion of strain numbers of fungal sequences, incorporating new species and strains as their genome sequences become available. This in turn will strengthen the accuracy and effectiveness of this optimised pipeline. Given the current dearth of these sequences in comparison to the prokaryotic kingdom, an increased effort needs to be made by the wider mycology research community to sequence new species and strains, assess mycobiome composition in different habitats and improve annotation of established and new fungal genomes. This will have the knock-on benefit of significantly improving our ability to model and analyse individual fungal species and communities with significant implications for future understanding of host-fungal interactions.

One of the main limitations of our strategy has been highlighted as the need for clarification of taxonomic similarity between different species. This can be only be improved by more species-specific genome annotation. Nevertheless, the innovation of species level fungal annotations has been demonstrated here. The accessibility and availability of the automated bioinformatic pipeline has now been improved through the use of a Nextflow pipeline, making this tool easier for widespread use. This work serves to assist in bringing advances in metagenomics tools and databases for analysis from the prokaryotic world to fungi and mycobiome. This will enable the community to drive deeper understanding of how the mycobiome affects human health and disease, including recent research in fungal associations with cancer.

## Data availability

The raw metagenome reads of the mock community, as well as oral and gut samples can be found PRJEB78538. The fungal catalogue can be found in paper reference here^36^ and tool is available at https://github.com/sysbiomelab.

## Acknowledgement

Many thanks to Professor Bernhard Hube (Hans Knoll Institute, Jena) for kindly supplying the *C. parapsilosis* and *P. jadinii* strains and to Professor Robin May (University of Birmingham) for the *Cryptococcus* species used in this study. SS was supported by Engineering and Physical Sciences Research Council (EPSRC), EP/S001301/1, Science for Life Laboratory, Swedish National Infrastructure for Computing at SNIC through Uppsala Multidisciplinary Centre for Advanced Computational Science (UPPMAX) under projects SNIC 2020-5-222, SNIC 2019/3-226, SNIC 2020/6-153. DLM and SS were supported by the Biotechnology Biological Sciences Research Council (BBSRC) BB/S016899/1,

## Author contributions

N.B, D.M. and S.S. conceived the project, drafted and edited the manuscript. N.B. performed sample preparation, extraction, experimental validation and optimisation, data preparation, extraction protocols for this paper. N.B. developed the bioinformatic pipeline, analysis and made all the figures included in this manuscript. S.L. provided intellectual input on design, statistical and analysis of the paper. M.A. and D.E. provided critical advice and input for bioinformatics pipeline. V.C.P. provided clinical samples for extraction. V.C.P., L.E., J.N. and M.U. provided critical feedback on the data and manuscript. All authors read, edited and reviewed the manuscript.

---

**Supplementary Table 1.** Strains that make up the mock community. Species, strains and references are provided.

**Supplementary Table 2.** Published primers for control, species specific for testing against mock community. Information based on sequence, melting temperature, concentration and reference provided.

**Supplementary Table 3.** Quantitative real-time PCR cyclic conditions.

**Supplementary Figure 1** In-vitro assessments of mock community. **A)** Assessment of DNA quantitation of mock community using ITS/ITS2 primers by qPCR. Green represents MGPS and orange represents PKS. Error bar represented by mean ± SD of biological triplicates. **B)** Normalised abundance of mock community using species-specific primers. Abundance normalised using inverse proportions of Ct value (Methods). Green represents MGPS and orange represents PKS.

**Supplementary Figure 2.** Evaluation of similarity, breadth and alignment coverage of fungal catalogue. **A)** Average nucleotide assessment between fungal species demonstrating nucleotide-level similarity. Pairwise analysis across fungal species with red indicating similarity and blue showing no similarity. **B)** Breadth coverage of mock community determined in both MGPS and PKS extraction to assess shot-gun sequence fragments covered by fungal catalogue. Each extraction has high coverage from fungal catalogue. Error bar represented by mean ± SD of biological triplicates. **C)** Percentage of fungal catalogue alignment to mock community from both extractions. PKS appears to have higher alignment than MGPS extraction. Error bar represented by mean ± SD of triplicates.

**Supplementary Figure 3.** Determine effect of extraction method on sequencing and alignment of unmapped reads. **A)** Percentage of mapped and unmapped mock community reads based on alignment against fungal catalogue across both chemical (MGPS) and kit-based (PKS) extraction methods. **B)** Read count of unmapped reads aligned using NCBI database revealing fungal species do not present in the fungal catalogue. Presence of other strains of species and similar sequences to mock community identified in unmapped reads.

**Supplementary Figure 4.** Assessment of the oral and gut patient samples. **A)** DNA concentration (ng/µl) of each healthy patient, including both oral and gut samples. **B)** Overall alignment of reads from oral and gut using fungal catalogue. **C)** Alignment of all reads to animal (Bos_tarus_UMDS), bacterial (hs_10_4_igc2), in-house fungal, human (GRCh38) and plant (A_thaliana_TAIR10) catalogues.

