## Supplementary figures and images for "Enhanced Computational and Experimental Approaches for Comparative Analysis of the Human Mycobiome"

### Supplementary Figure 1

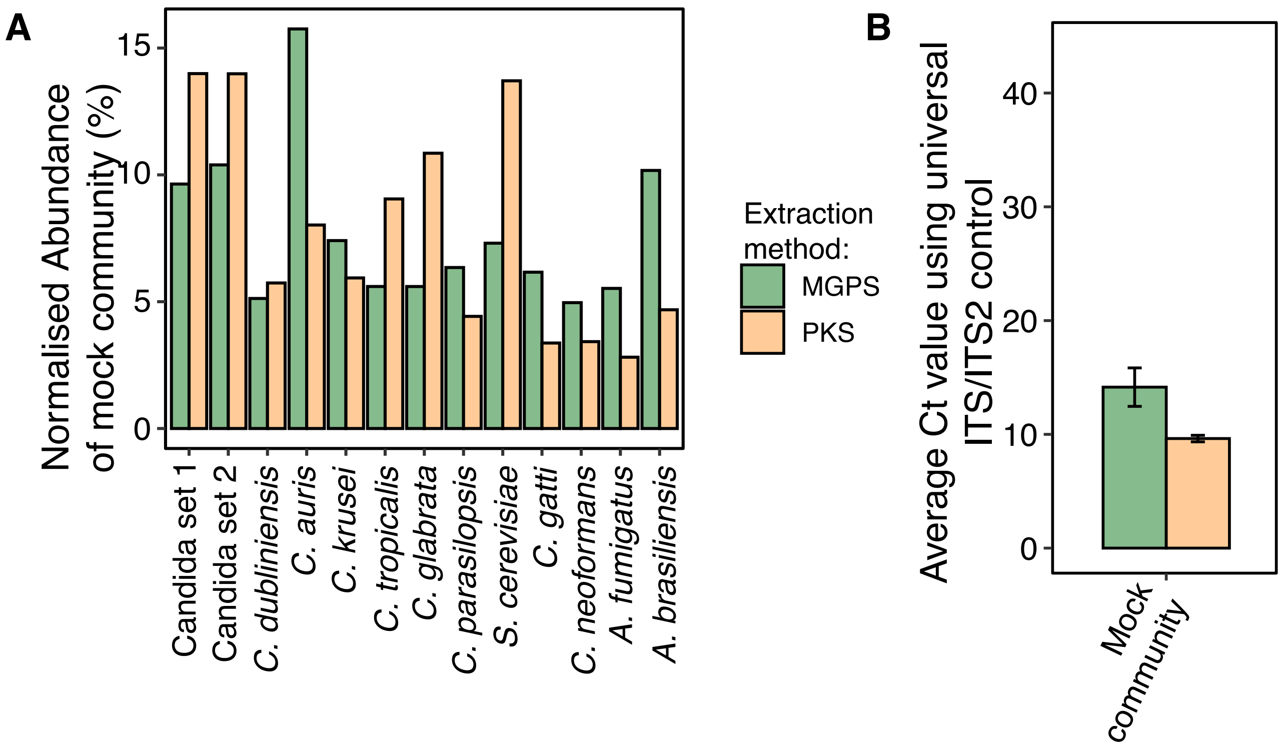

### Supplementary Figure 2

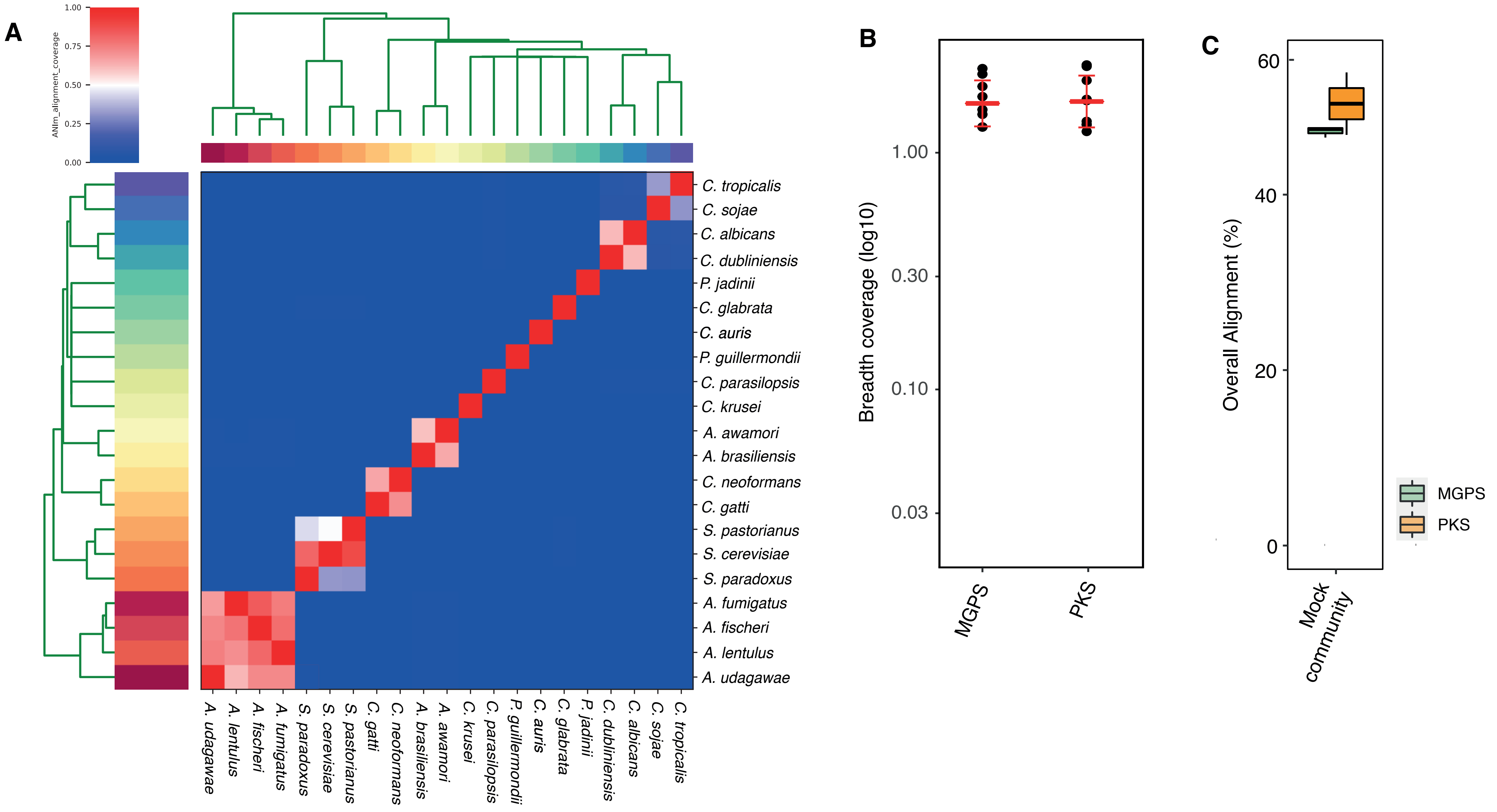

### Supplementary Figure 3

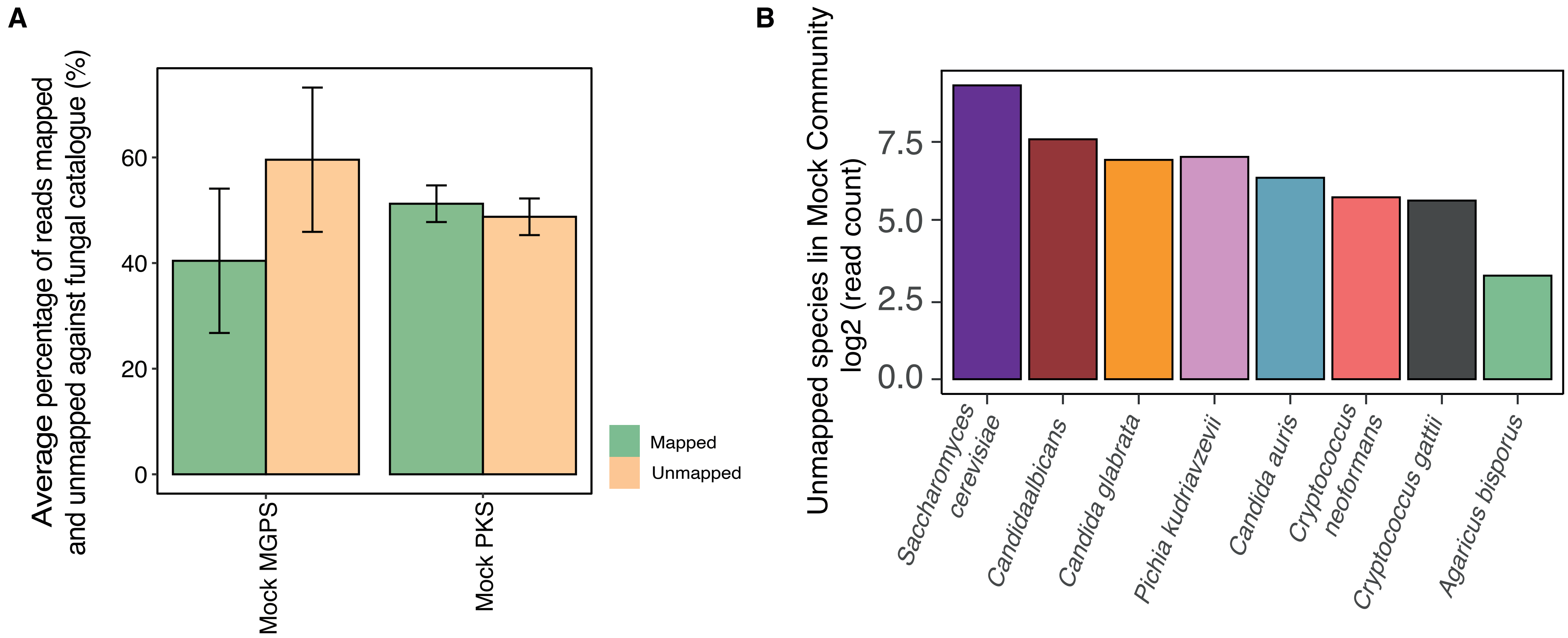

### Supplementary Figure 4

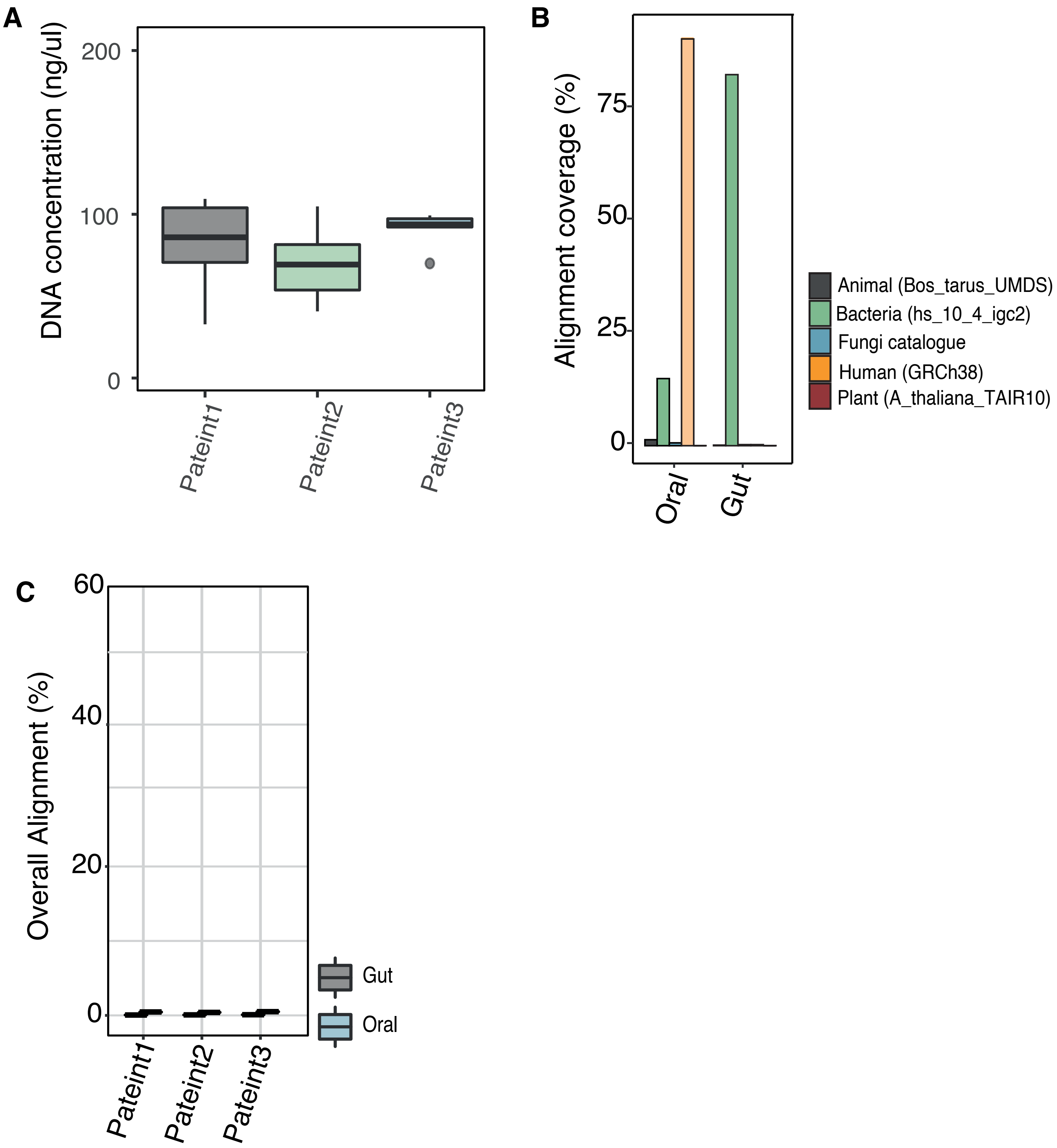
